# Scalable phylodynamic inference of mutational and epistatic fitness effects with multi-feature birth-death models

**DOI:** 10.64898/2026.10.19.760109

**Authors:** Shi Cen, Lenora M. Kepler, Cristina Lanzas, Brian J. Reich, David A. Rasmussen

**Affiliations:** Bioinformatics Research Center, North Carolina State University, Raleigh, North Carolina, USA; Department of Statistics, North Carolina State University, Raleigh, North Carolina, USA; Department of Population Health and Pathobiology, North Carolina State University, Raleigh, North Carolina, USA; Department of Entomology and Plant Pathology, North Carolina State University, Raleigh, North Carolina, USA

**Keywords:** Genotype-to-fitness map, phylodynamics, multi-feature birth-death model, fitness landscape, epistasis, Hamiltonian Monte Carlo, antimicrobial resistance, QRDR mutations

## Abstract

Fitness is a complex phenotype determined by many different genomic features, their interactions with each other and the environment. For rapidly evolving microbes like viruses and bacteria, phylodynamic multi-feature birth-death (MFBD) models provide a way to infer the fitness of lineages with different genotypes from reconstructed phylogenies. However, existing phylodynamic methods assume very simple additive or multiplicative genotype-to-fitness (G-to-F) mappings that typically ignore epistatic interactions among features. We developed a gradient-based Bayesian MFBD framework for estimating the fitness effects of multiple features while allowing for both additive and epistatic G-to-F maps. The framework uses Hamiltonian Monte Carlo for posterior inference and incorporates shrinkage priors to regularize weakly supported fitness effects without cross-validation. In simulations, Bayesian inference performs well at recovering additive fitness effects, with accuracy and precision improving consistently with increasing tree size. While epistatic effects are more difficult to estimate because they depend on the co-occurrence of mutations, shrinkage improves estimation when interaction effects are weak or sparsely supported. Simulations with misspecified additive models show that ignoring moderate or strong epistasis can bias estimates of individual mutational effects. We applied the framework to a time-calibrated *Escherichia coli* ST131 phylogeny with 1763 samples and estimated fitness effects for several mutations in the quinolone resistance-determining region (QRDR). These results show that Bayesian phylodynamic inference can be used to learn the genomic determinants of fitness while quantifying uncertainty in mutational and epistatic fitness effects.

## Introduction

Estimating the fitness effects of mutations and other genomic features is central to understanding the genetic basis of adaptation. These fitness effects determine whether a lineage is more likely to grow, persist, and spread, and they therefore shape the evolutionary trajectories available to a population (Greenbury *et al*., 2022; Aguilar-Rodríguez *et al*., 2017). In microbial pathogens, fitness-effect estimation has direct epidemiological relevance because genetic changes affecting antimicrobial resistance, immune escape, replication, persistence, or transmission can alter the population-level success of a lineage (Kepler *et al*., 2021). Genotype-to-fitness (G-to-F) maps provide a useful conceptual framework for this problem by describing how genetic variation translates into differences in evolutionary success or fitness (Visser and Krug, 2014). G-to-F maps can be represented as fitness landscapes, where genotypes occupy points in a high-dimensional sequence space and fitness defines a surface over that space (Wright, 1980; Bank, 2022). From an inference perspective, the practical goal is to estimate which genomic features contribute to fitness variation, how large those effects are, and how much uncertainty remains around them.

Estimating fitness effects is challenging because the effect of a mutation can depend on the genetic background and changing environments in which it occurs. Epistasis occurs when the phenotypic or fitness effect of one mutation is modified by other mutations, creating fitness landscapes with multiple peaks and valleys (Bank, 2022; Johnson *et al*., 2023). Idiosyncratic epistasis arises from interactions among specific loci, including higher-order interactions involving two or more background mutations, and can make individual mutational effects difficult to identify (Sailer and Harms, 2017). These dependencies make it harder to attribute differences in lineage growth or transmission to individual mutations.

High-throughput mutagenesis experiments in controlled environments are usually required to learn the properties of G-to-F maps (Zhou *et al*., 2022). These experiments directly measure many engineered genotypes and can characterize local landscape structure with high precision. Experimentally measured fitness, however, depends on the assay environment and may not be representative of fitness in real-world environments (Blanquart and Bataillon, 2016). Complete combinatorial libraries also become infeasible as sequence length or interaction order increases (Fowler and Fields, 2014). Phylodynamic methods, on the other hand, provide a complementary approach and have been increasingly used to learn the genomic determinants of pathogen fitness in real-world settings (Kepler *et al*., 2021; Lefrancq *et al*., 2025; Helekal *et al*., 2023). Phylogenies record the branching events of lineages, allowing differences in lineage growth or transmission to be connected to their genetic features (Volz *et al*., 2013).

Multi-type birth-death (MTBD) models formalize the connection between genotype and fitness by assigning lineages to types with distinct birth, death, or sampling rates (Stadler and Bonhoeffer, 2013; Barido-Sottani *et al*., 2020). The estimated type-specific birth and death rates can then be used to impute the growth rate or fitness of each lineage. These models are increasingly used to quantify the fitness effects of individual mutations or other genomic features, including the transmission cost of HIV drug-resistance mutations (Kühnert *et al*., 2018) and multidrug resistance in Mycobacterium tuberculosis (Pečerska *et al*., 2021). Rasmussen and Stadler (2019) extended MTBD models to cases where multiple sites in a sequence jointly determine the fitness of a lineage as they evolve. More recently, these *multi-feature* birth-death (MFBD) models have been extended to allow for a large number of genomic features to determine the fitness of lineages and for their individual contributions to population-level fitness to be estimated from a reconstructed phylogeny (Kepler *et al*., 2021, 2024). These developments make it possible to learn the genomic determinants of pathogen fitness in the epidemiological settings where selection occurs.

While MFBD models offer a powerful tool for learning the mapping between multiple features and the fitness, existing methods suffer from several limitations. First, the power to estimate the fitness effects of individual features largely depends on the phylogenetic information available to estimate them (i.e. having sampled a sufficient number of lineages with and without a given feature). This problem becomes more severe when epistatic interactions are included because only a small subset of all possible mutational combinations will occur in sampled lineages. Second, maximum likelihood MFBD approaches (Kepler *et al*., 2021, 2024) obtain point estimates of fitness effects but do not allow for full uncertainty quantification, which may be especially important for rare features or features with small fitness effects. Third, when estimating idiosyncratic epistasic interactions between features that co-occur at low prevalence, regularization is needed to avoid overfitting. However, choosing an optimal regularization factor usually requires running cross-validation, which is computationally expensive. These limitations have restricted the application of MFBD models to considering relatively small sets of features with additive or multiplicative effects on fitness.

In this work, we develop a gradient-based Bayesian MFBD inference framework for estimating additive and idiosyncratic epistatic G-to-F maps from phylogenies, allowing for more efficient and faster high-dimensional inference on mutational fitness effects using Hamiltonian Monte Carlo with uncertainty quantification of the fitness effect estimates. We evaluate the framework using simulated phylogenies with four or ten genetic features and varying numbers and strengths of pairwise interactions. Shrinkage priors are applied with data-informed hyperparameters without requiring cross-validation when estimating individual and epistatic fitness effects. The simulations compare Bayesian estimates with maximum likelihood estimates and quantify how accuracy depends on tree size, feature collinearity, mutation co-occurrence, and prior choice. Finally, we apply this gradient-based MFBD framework to a time-calibrated phylogeny of *E. Coli* ST131 and estimate additive, pairwise, and higher-order effects among quinolone resistance-determining region (QRDR) mutations, which allows us to compare the estimated fitness effect against previous lab-based results (Ben Zakour *et al*., 2016; Baker *et al*., 2013; Marcusson *et al*., 2009). Together, this framework provides uncertainty-aware inference of the genetic determinants of fitness in naturally circulating populations.

## Methods

### Overview of the multi-type birth-death model

MTBD models extend birth-death-sampling models by allowing lineages to belong to discrete types with type-specific epidemiological or evolutionary rates (Maddison *et al*., 2007; Stadler and Bonhoeffer, 2013; Barido-Sottani *et al*., 2020). In pathogen phylodynamics, birth events are usually interpreted as transmission, death events as removal or loss of infectiousness, and sampling events as the observation of infected hosts or pathogen genomes (Figure 1A)(Volz *et al*., 2013). The probability of an observed time-calibrated phylogeny is then determined by the branching times, sampling times, lineage types, and the rates assigned to each type. In this study, each type corresponds to a genotype or feature combination, and the type-specific birth rate is interpreted as a population-level fitness parameter. This links genetic features directly to differences in lineage growth or transmission fitness.

**Figure 1:**
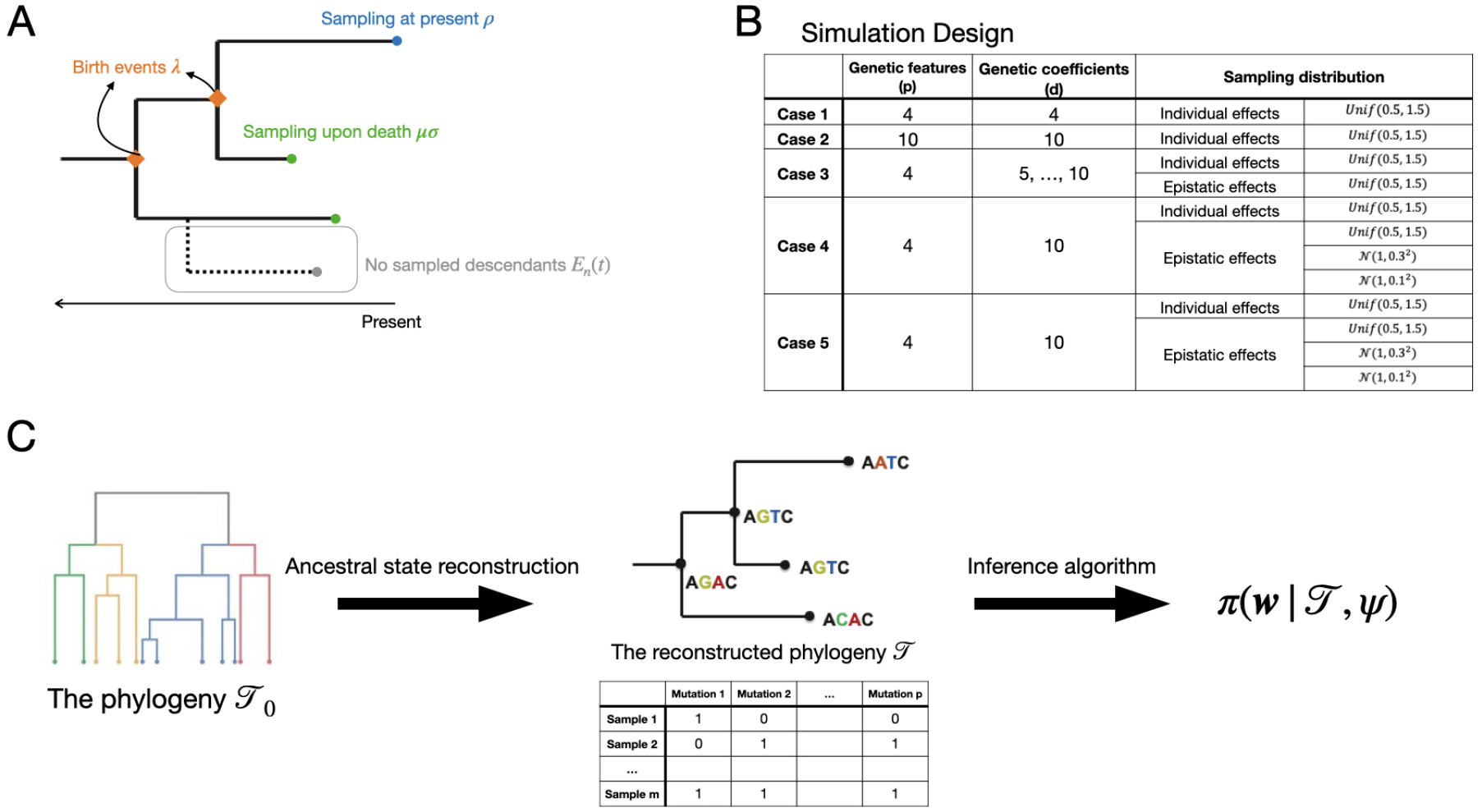
(A) Schematic of the birth-death-sampling process. The phylogenetic likelihood computes the probability density of the tree evolving as observed under this process. Birth events occur at rate λ, sampling upon removal occurs at rate *µ*σ, and extant lineages are sampled at the present with probability *ρ*. Unsampled descendant lineages are accounted for through *E*_*n*_ (*t*), the probability a lineage has no sampled descendants. (B) Simulation scenarios used to evaluate inference performance across additive and epistatic fitness models with different genome sizes, numbers of inferred features, and prior distributions for individual and epistatic effects. (C) Analysis workflow for empirical datasets. A time-calibrated phylogeny *T*_0_ and tip feature states are used to reconstruct ancestral states, producing a feature-annotated phylogeny *T* and character matrix for Bayesian inference of the posterior distribution *π* (**w** |*T*, ψ) . Note that the state feature is a probability valued between 0 and 1.

### Likelihood of multi-feature birth-death model

Under a MFBD model, the likelihood of the phylogeny can be calculated as previously described in Barido-Sottani *et al*. (2018) and Kepler *et al*. (2024). The likelihood function is given by:

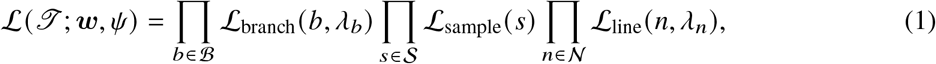

where *T* is the observed phylogeny, ψ = (σ, *µ, ρ*) is the set of the fixed parameters (σ-Sampling probability upon removal/death, *µ*-Removal/Death rate, *ρ*-Sampling proportion at present) which are assumed to be known.

The likelihood (1) can therefore be factorized into a product of the likelihood of each branching event (*ℒ* _branch_), sampling event (*ℒ* _sample_) and lineage evolving as observed (*ℒ* _line_). These components of the likelihood are computed as:

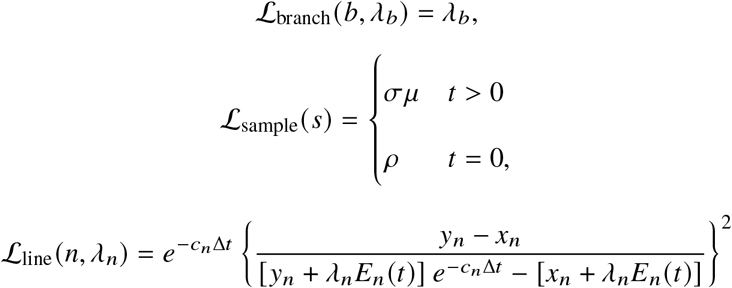

with:

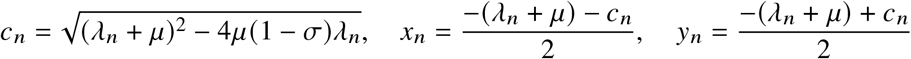

The probability *E*_*n*_ (*t*) that a lineage at time *t* in the past produced no sampled descendants is given by:

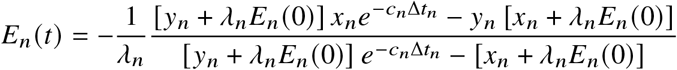

Here, we let the probability of lineage not being sampled at the present *E*_*n*_ (0) = 1 − *ρ*.

In this framework, each genotype defines a lineage type, and the genotype-specific fitness value determines the corresponding birth rate λ. The phylogenetic likelihood therefore links the observed branching and sampling events to the latent genotype-to-fitness map. This likelihood serves as the basis for both Bayesian posterior inference and maximum likelihood estimation.

Because lineages and events are conditionally independent given the model parameters, the phylogenetic likelihood can be evaluated as a series of matrix-vector operations across all tree components. We implemented these operations in JAX for efficient vectorized computation (Bradbury *et al*., 2018). We also use JAX’s automatic differentiation methods to compute the likelihood gradients required for Hamiltonian Monte Carlo, avoiding the need for analytic derivatives.

### Statistical inference

GivenS a fixed phylogenetic tree, we assume that the features ***x*** of all ancestral lineages can be reconstructed from the observed genotypes at the tip of the tree. We used Bayesian inference to estimate the fitness effects of individual mutations 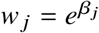 and pairwise epistasis 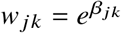 the observed and reconstructed genotypes *X*_*m*×*p*_ in the phylogeny, where *m* = | *ℬ* | + |*N* | is the total number of branching events and lineages. We compare estimation performance under three priors for the fitness effects:

- *w* _*j*_, *w* _*jk*_ *∼* Unif(0.0, 2.0)
- *w* _*j*_, *w* _*jk*_ *∼ N* (1, *τ*^2^)1(*w* _*j*_ > 0), *τ*^2^ *∼* InvGamma(1, 1)
- *w* _*j*_, *w* _*jk*_ *∼* Laplace(1, *τ*)1(*w* _*j*_ > 0), *τ*^2^ *∼* InvGamma(1, 1).

The uniform prior is used as a weakly informative baseline, whereas the normal and Laplace priors place greater prior mass near one and therefore provide regularization against overfitting. For the shrinkage priors, the variance, and thus the strength of shrinkage, is inferred from the phylogeny. The hierarchical Bayesian model for the additive fitness landscape is specified as follows:

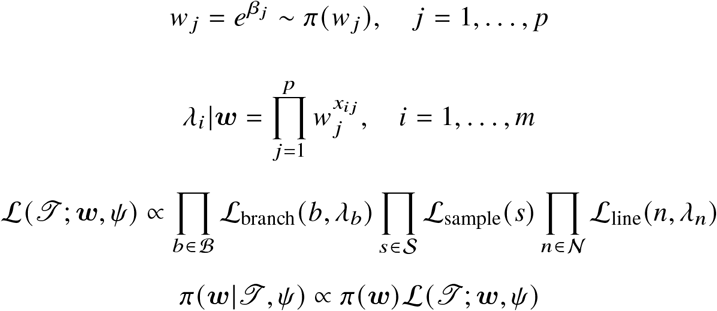

Posterior inference was performed using Markov chain Monte Carlo. For each parameter, the posterior median was reported as the point estimate because it is invariant under monotone transformations and is robust to posterior skewness. Equal-tailed 95% credible intervals were computed from the posterior samples.

For comparison, we also performed inference under the maximum likelihood estimator (MLE)

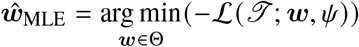

as a frequentist benchmark. The asymptotic variance of the MLE is given by the Cramer-Rao lower bound

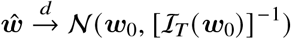

where *ℐ*_*T*_ (***w***_0_) is the total Fisher information matrix and can be approximated by the observed information matrix 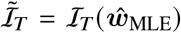. Wald-type 95% confidence intervals were constructed using the asymptotic variance of the MLE. We compared Bayesian and maximum likelihood estimates using empirical coverage, mean absolute error (MAE), interval width and correlation coefficients between the true and estimated values.

### Simulation Study

We evaluated the proposed inference framework using simulated phylogenies generated under a birth-death-sampling process. Trees were simulated with the Gillespie stochastic algorithm (Gillespie, 2007) starting with a single infected individual. For each lineage, fitness was specified as a linear function of the genotype-specific additive and, when applicable, epistatic effects. For lineage *i*, we denote its genotype ***x***_*i*_ = (*x*_*i*1_, · · ·, *x*_*i p*_)^⊺^ ∈ {0, 1}^*p*^ where *p* is the number of genetic features. In the additive model, the latent fitness layer is defined as 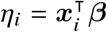, where *β*_*i*_ is the fitness effect of a given feature on the log-scale. Fitness effects are therefore assumed to be multiplicative on a linear scale and constrained to be positive.

Lineage fitness λ_*i*_ is obtained through the mapping function 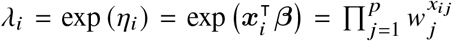 where *w* _*j*_ = exp( *β* _*j*_ ) are the multiplicative fitness effects.

To evaluate statistical performance under different levels of complexity in the genotype-to-fitness map, we varied the number of genetic features *p* and the number of included interaction terms *q*, giving *d* = *p* + *q* genetic coefficients. In the additive scenarios, we considered alignments with either four or ten genetic features (Case 1 and 2). To study pairwise epistasis, interaction terms were constructed from first-order products among genetic features (Case 3 to 5). Individual mutational effects *w* _*j*_ were sampled from the Unif(0.5, 1.5) distribution, and the corresponding epistatic effects *w* _*jk*_ were sampled from uniform or normal distributions with a variance term that controls the magnitude of epistatic interactions. Larger variances therefore tend to generate stronger interactions, whereas smaller values generate weaker interactions (Case 4). To test whether ignoring epistasis biases fitness effect estimates, we analyzed the Case 4 datasets under a reduced inference model that excluded epistatic terms (Case 5). Specifically, we have the following simulation settings (Figure 1B):

- Case 1: *p* = 4, *q* = 0, *d* = 4, *w* _*j*_ *∼* Unif(0.5, 1.5)
- Case 2: *p* = 10, *q* = 0, *d* = 10, *w* _*j*_ *∼* Unif(0.5, 1.5)
- Case 3: *p* = 4, *q* = 1, …, 6, *d* = 4 + *q* with *w* _*j*_ *∼* Unif(0.5, 1.5), *w* _*jk*_ *∼* Unif(0.5, 1.5)
- Case 4: *p* = 4, *q* = 6, *d* = 4 + 6 = 10
  - Strong epistasis: *w* _*j*_ *∼* Unif(0.5, 1.5), *w* _*jk*_ *∼* Unif(0.5, 1.5)
  - Moderate epistasis: *w* _*j*_ *∼* Unif(0.5, 1.5), *w* _*jk*_ *∼* N (1, 0.3^2^)
  - Weak epistasis: *w* _*j*_ *∼* Unif(0.5, 1.5), *w* _*jk*_ *∼* N (1, 0.1^2^)
- Case 5: *p* = 4. The true model has *q*_true_ = 6 and *d*_true_ = 10, whereas the reduced fitted model has *q*_fit_ = 0 and *d*_true_ = 4. The true effects follow the same regimes as Case 4:
  - Strong epistasis: *w* _*j*_ *∼* Unif(0.5, 1.5), *w* _*jk*_ *∼* Unif(0.5, 1.5)
  - Moderate epistasis: *w* _*j*_ *∼* Unif(0.5, 1.5), *w* _*jk*_ *∼ N* (1, 0.3^2^)
  - Weak epistasis: *w* _*j*_ *∼* Unif(0.5, 1.5), *w* _*jk*_ *∼ N* (1, 0.1^2^)

We additionally evaluated the interaction-strength comparison with ten genetic features (*p* = 10), including all 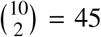 possible pairwise interaction terms (*q* = 45), giving *d* = *p* + *q* = 55 genetic coefficients. As in Case 4, main-effect multipliers follow *w* _*j*_ *∼* Unif(0.5, 1.5), while pairwise interaction multipliers follow *w* _*jk*_ *∼* Unif(0.5, 1.5) under strong epistasis, *w* _*jk*_ *∼* N (1, 0.3^2^) under moderate epistasis, and *w* _*jk*_ *∼* N (1, 0.1^2^) under weak epistasis. We jointly estimated the main and pairwise interaction effects under each of the three prior distributions and summarized their recovery separately (Table 7).

## Regression Analysis

To identify factors associated with estimation accuracy, we regressed mean absolute error (MAE) against simulation-derived features from each phylogeny. We denote phylogeny size by |*T* |, defined as the total number of tips. We use the condition number to measure the overall multicollinearity of the feature matrix, calculated as 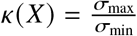 where σ_max_ and σ_min_ are the largest and smallest singular values of *X*, respectively. The number of occurrences or co-occurrences, *N*_(co-)occur_, is defined as the total number of lineages carrying a given mutation or, for idiosyncratic epistasis, a given mutation pair.

### Computational implementation

Our Bayesian inference framework, BayesMFBD, is implemented in Python and builds on several existing scientific computing libraries. We performed Bayesian inference via Markov chain Monte Carlo (MCMC) implemented in NumPyro (Phan *et al*., 2019). Posterior samples were drawn using the No-U-Turn Sampler (NUTS, (Hoffman and Gelman, 2011)), an adaptive extension of Hamiltonian Monte Carlo (HMC) that eliminates the need to manually tune the number of leapfrog steps. We ran one independent chain each with 1000 warm-up (tuning) steps followed by 5000 sampling steps. Convergence was assessed using the potential scale reduction factor (*R* < 1.01 for all parameters) and effective sample size (ESS > 400 per chain). All sampling was performed on JAX-accelerated hardware via NumPyro’s XLA backend.

### Analysis workflow

The same workflow was used to prepare simulated and empirical datasets for phylodynamic inference (Figure 1C). The required inputs are a rooted time-calibrated phylogeny *T*_0_ and a character matrix giving the observed states of the fitness-determining features at the tips. For simulated datasets, the phylogeny and feature states were generated directly under the specified birth-death-sampling model. For the ST131 analysis, the time-calibrated phylogeny and QRDR genotypes were obtained from the genomic data as described in (Kepler *et al*., 2024).

A typical workflow consists of the following steps:

- **Input files:** Users provide a rooted time-calibrated phylogeny, which can be reconstructed in BEAST2 (Bouckaert *et al*., 2014) or IQTree(Wong *et al*., 2026), and a feature matrix containing the observed states of candidate fitness-determining features at the sampled tips. These features may represent mutations, genes, phenotypes, or other discrete lineage attributes.
- **Ancestral-state reconstruction:** If internal-node states are not already available, ancestral states are reconstructed on the phylogeny. This produces a feature-annotated tree in which each branch is assigned an inferred feature state. The inferred states can be binary or probabilistic values between 0 and 1.
- **Model specification:** Users specify the fitness model by choosing whether to include additive effects alone or additional interaction terms for candidate epistatic effects. Prior distributions and other fixed MFBD parameters are also specified before inference.
- **Tree vectorization:** The pipeline converts the annotated phylogeny into arrays of branching events, sampling events, lineage intervals, and feature states. These arrays are used to evaluate the multi-feature birth-death likelihood.
- **Posterior inference:** The pipeline estimates posterior distributions for additive and epistatic fitness effects using the specified Bayesian model and MCMC settings.
- **Outputs:** The main outputs are posterior summaries for each fitness effect, including point estimates, credible intervals, and diagnostic quantities indicating whether each feature is sufficiently supported by the phylogenetic data.

## Results

### Additive genotype-to-fitness maps

We first evaluated the performance of our estimator with the additive fitness model (Case 1 and 2) in a low-dimensional setting with four genetic features. For each simulation replicate, we estimated the four individual mutational fitness effects and summarized performance using coverage, mean absolute error (MAE), and credible or confidence interval width. In addition to these overall measures of performance, to better understand how features of the dataset and prior affect performance, we regressed MAE against genotype-derived features from each phylogeny. The association between MAE and other genotype-derived features is modeled as:

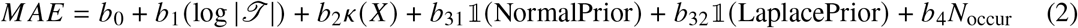

Bayesian inference and maximum likelihood estimation produce broadly comparable performance in this setting, with similar coverage, MAE, and interval width (Table 1). Under the uniform prior, estimated credible intervals generally capture the true effects, and the effects of mutations that occurred more frequently in the alignment tend to be estimated more accurately (Figure 2A). This pattern is expected because recurrent mutations provide more phylogenetic information about their contribution to lineage fitness. Estimated fitness effects tend to be biased upwards for highly deleterious mutations, such that they are estimated to be less deleterious than they actually are. Across the three Bayesian priors, estimation accuracy is similar, suggesting that shrinkage provides limited benefit when the number of genetic features is small and the additive model is correctly specified (Table 1, 2). Tree size is the strongest determinant of accuracy (Table 2, Figure 2B). MAE decreases significantly as tree size increased, indicating that larger phylogenies provide more information for separating mutation-specific fitness effects. By contrast, the condition number of the feature matrix is not significantly associated with MAE in this low-dimensional setting (Table 2, Figure 2C), suggesting that moderate feature correlation does not substantially impair inference when only four additive effects are estimated.

**Table 1:** Estimation performance for the four-feature additive model.

| Priors | Coverage |  | MAE |  | CI width |  | Correlation |  |
| --- | --- | --- | --- | --- | --- | --- | --- | --- |
|  | MCMC | MLE | MCMC | MLE | MCMC | MLE | MCMC | MLE |
| Uniform | 0.873 |  | 0.072 |  | 0.315 |  | 0.939 |  |
| Normal | 0.883 | 0.910 | 0.072 | 0.071 | 0.315 | 0.357 | 0.948 | 0.935 |
| Laplace | 0.870 |  | 0.074 |  | 0.313 |  | 0.941 |  |

**Table 2:** Regression of MAE on tree size, feature collinearity, prior distribution and number of independent occurrence of mutations for the four-feature additive model.

|  | Estimate | Std error | t value | P value |
| --- | --- | --- | --- | --- |
| Intercept | 0.215 | 0.035 | 6.172 | < 0.001 |
| $\log \vert \mathcal{T}\vert $ | -0.019 | 0.005 | -3.844 | < 0.001 |
| $\kappa(X)$ | -0.006 | 0.006 | 0.899 | 0.369 |
| Prior (Normal) | 0.002 | 0.005 | 0.495 | 0.621 |
| Prior (Laplace) | 0.005 | 0.005 | 1.022 | 0.307 |
| $N_{\text{occur}}$ | < 0.001 | < 0.001 | -0.769 | 0.443 |

**Figure 2:**
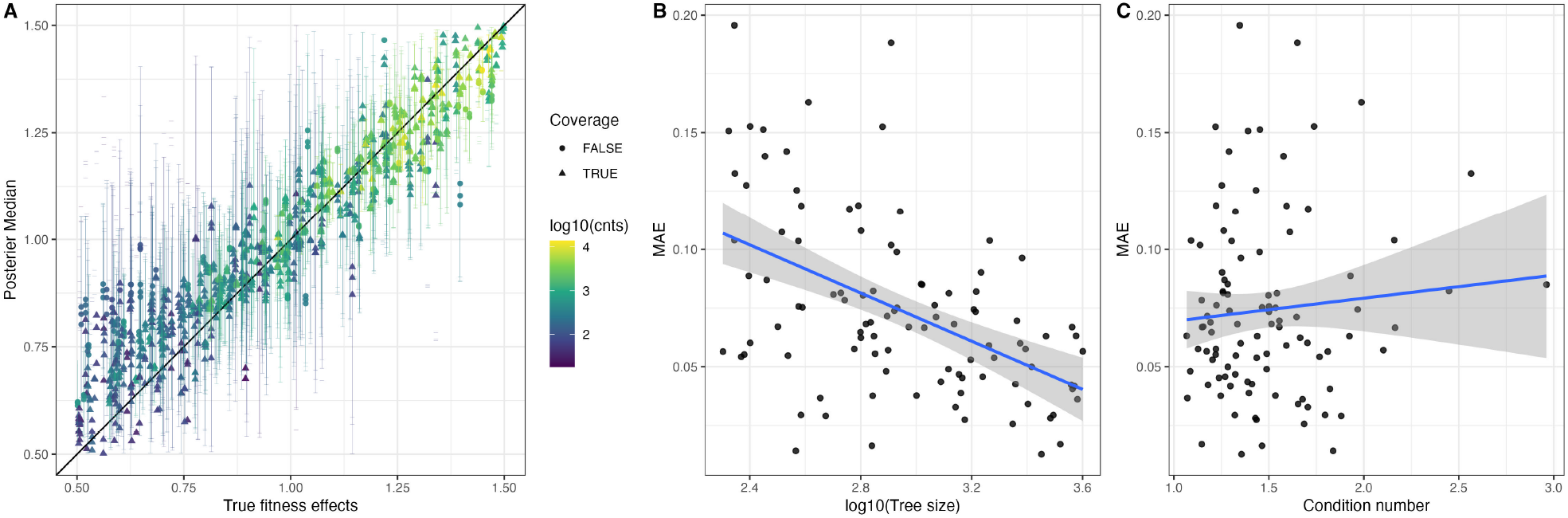
Estimation performance for the four-feature additive fitness model using a uniform prior. (A) Posterior median estimates of mutational fitness effects plotted against the true simulated fitness effects. Points are colored by the log-transformed number of independent occurrences of each mutation in the simulated alignment. Triangles indicate that the true value falls within the 95% credible interval, and circles indicate that it does not. Vertical error bars represent 95% credible intervals, and the diagonal line denotes perfect agreement between estimated and true effects. (B) Mean absolute error (MAE) plotted against the log-transformed tree size. A regression line was fitted with a shaded confidence band. (C) MAE plotted against the condition number of the genotype feature matrix. The line and shaded band are as in B.

We next considered an additive model with ten genetic features. The overall average MAE in the ten-feature setting is larger than in the four-feature setting, indicating that estimation becomes more difficult as the number of fitness-effecting features increases relative to the amount of phylogenetic information (Tables 3 and 4). Tree size remains a strong determinant of performance and is associated with significantly lower MAE (Figure 3A, Table 4). Neither collinearity nor prior choice materially changes inferential performance (Figure 3B, Table 4). Accuracy thus scales predictably with phylogeny size across simulations regardless of the number of features, establishing the foundation for assessing the harder problem of pairwise epistatic inference.

**Table 3:** Estimation performance for the ten-feature additive model.

| Priors | Coverage |  | MAE |  | CI width |  | Correlation |  |
| --- | --- | --- | --- | --- | --- | --- | --- | --- |
|  | MCMC | MLE | MCMC | MLE | MCMC | MLE | MCMC | MLE |
| Uniform | 0.927 |  | 0.105 |  | 0.021 |  | 0.828 |  |
| Normal | 0.926 | 0.956 | 0.106 | 0.110 | 0.021 | 0.027 | 0.860 | 0.769 |
| Laplace | 0.917 |  | 0.108 |  | 0.022 |  | 0.849 |  |

**Table 4:** Regression of MAE on tree size, feature collinearity, and prior distribution for the ten-feature additive model.

|  | Estimate | Std error | t value | P value |
| --- | --- | --- | --- | --- |
| Intercept | 0.417 | 0.017 | 25.012 | < 0.001 |
| $\log \vert \mathcal{T}\vert $ | -0.044 | 0.002 | -20.183 | < 0.001 |
| $\kappa(X)$ | -0.005 | 0.003 | -1.635 | 0.103 |
| Prior (Normal) | 0.001 | 0.005 | 0.152 | 0.879 |
| Prior (Laplace) | 0.003 | 0.005 | 0.615 | 0.539 |

**Figure 3:**
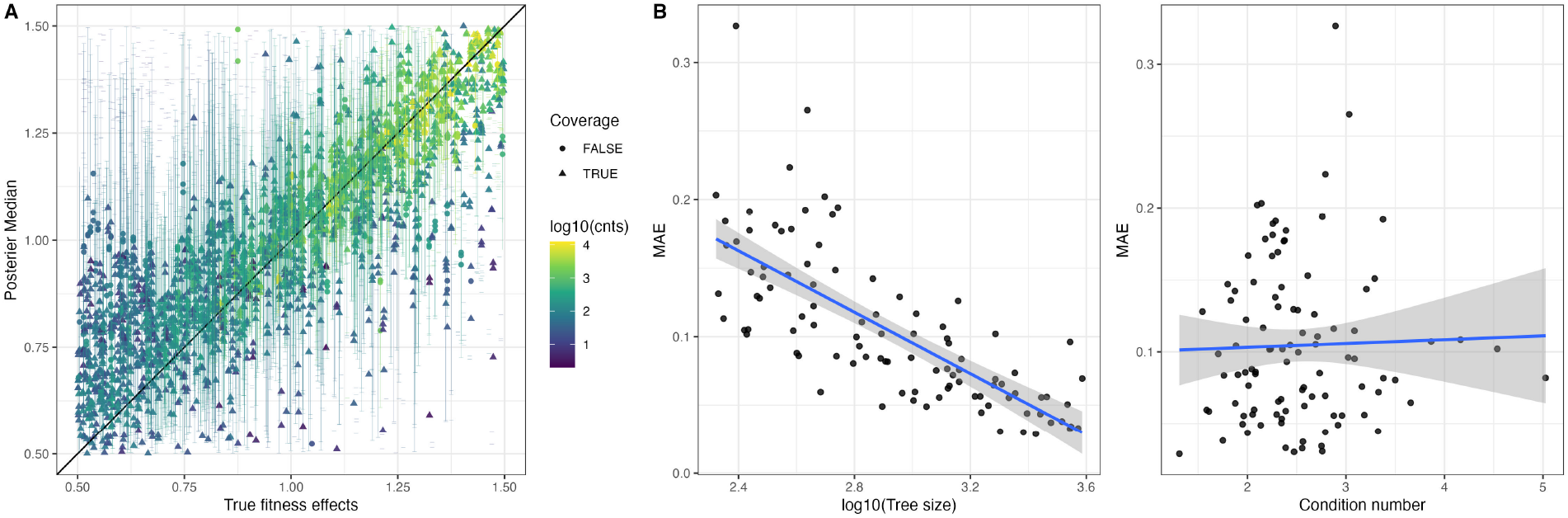
Estimation performance for the ten-feature additive fitness model. (A) Posterior median estimates of mutational fitness effects plotted against the true simulated fitness effects. Points are colored by the log-transformed number of occurrences of each mutation in the simulated alignment. Triangles indicate that the true value falls within the 95% credible interval, and circles indicate that it does not. Vertical error bars represent 95% credible intervals, and the diagonal line denotes perfect agreement between estimated and true effects. (B) Mean absolute error (MAE) plotted against the log-transformed tree size. A regression line was fitted with shaded confidence band (C) MAE plotted against the condition number of the genotype feature matrix. The line and shaded band are as in B.

### Pairwise epistasis in genotype-to-fitness maps

We next evaluated inference performance when the true genotype-to-fitness map includes pairwise epistatic interactions between features (Cases 3 and 4). To assess the effect of model complexity, we constructed six simulation subcases by adding different numbers of first-order interaction terms (from one to six). The association between MAE and other genotype-derived features is modeled as:

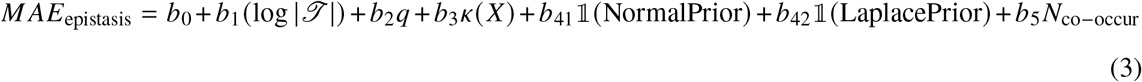

In general, accurately inferring epistatic interactions is more difficult than inferring the fitness effects of individual features, as a pairwise interaction can be informed only by lineages carrying the corresponding pair of mutations, and such combinations occur less frequently than individual mutations. As expected, the accuracy at which epispastic coefficients are estimated improves significantly with the the number of mutation co-occurrences (Table 5). Increasing the number of epistatic interactions does not significantly decrease overall MAE (Table 5).

**Table 5:** Regression of MAE on tree size, the number of epistasic pairs, feature collinearity, prior distribution, and number of co-occurrences of mutations.

|  | Estimate | Std error | t value | P value |
| --- | --- | --- | --- | --- |
| Intercept | 0.5667 | 0.06932 | 8.175 | < 0.001 |
| $\log \vert \mathcal{T}\vert $ | -0.0523 | 0.009580 | -5.463 | < 0.001 |
| $q$ | $7.954 \times 10^{-5}$ | $3.335 \times 10^{-3}$ | 0.024 | 0.9810 |
| $\kappa(X)$ | -0.0154 | 0.01372 | -1.124 | 0.2612 |
| Prior (Normal) | -0.0234 | 0.01180 | -1.982 | 0.0478 |
| prior (Laplace) | -0.0186 | 0.01180 | -1.575 | 0.1156 |
| $N_{\text{co-occur}}$ | $-3.326 \times 10^{-5}$ | $1.656 \times 10^{-5}$ | -2.009 | 0.0449 |

Applying a moderate level of shrinkage, i.e., a Normal prior, on the epistasis coefficients marginally decreases the absolute error of the estimates, but applying a strong shrinkage prior does not improve accuracy. This result suggests that the MFBD framework can accommodate a moderate number of interaction terms, provided that the phylogeny contains sufficient numbers of lineages with the relevant genotypes. Tree size remains a significant determinant of accuracy, with larger phylogenies producing more accurate estimates (Table 5). Prior choice had a modest effect. Relative to the uniform prior, the normal prior only provides a marginally significant improvement in accuracy, whereas the Laplace prior does not show a statistically significant improvement.

Next, we examined how the strength of epistasis (i.e. the magnitude of the epistatic interactions) impacts inference across priors under settings with four or ten genetic features. Across weak, moderate, and strong epistatic regimes, epistatic effects are consistently more difficult to estimate than individual mutational effects in both 4-feature and 10-feature models (Figure 4), with larger MAE and wider intervals (Tables 6, 7). Shrinkage priors provide clear advantages when epistatic effects are present, particularly under weak epistasis. In this regime, the normal and Laplace priors achieve comparable performance for individual mutational effects but substantially reduces the MAE and credible interval width for epistatic effects compared to the uniform prior. These results suggest that regularization is especially useful for estimating sparse or weakly supported interaction effects in genotype-to-fitness maps.

**Table 6:** Estimation performance with weak, moderate and strong epistatic interactions among four genetic features under different priors.

|  |  | Priors | Coverage | MAE | CI width | Correlation |
| --- | --- | --- | --- | --- | --- | --- |
| Strong epistasis<br>$w_j \sim \text{Unif}(0.5, 1.5)$<br>$w_{jk} \sim \text{Unif}(0.5, 1.5)$ | Individual effects | Uniform | 0.938 | 0.108 | 0.556 | 0.885 |
|  |  | Normal | 0.933 | 0.107 | 0.537 | 0.890 |
|  |  | Laplace | 0.935 | 0.108 | 0.523 | 0.889 |
|  | Epistatic effects | Uniform | 0.943 | 0.212 | 1.145 | 0.587 |
|  |  | Normal | 0.943 | 0.196 | 1.090 | 0.608 |
|  |  | Laplace | 0.943 | 0.193 | 1.067 | 0.610 |
| Moderate epistasis<br>$w_j \sim \text{Unif}(0.5, 1.5)$,<br>$w_{jk} \sim \text{Normal}(1, 0.3^2)$ | Individual effects | Uniform | 0.943 | 0.105 | 0.575 | 0.889 |
|  |  | Normal | 0.940 | 0.101 | 0.556 | 0.896 |
|  |  | Laplace | 0.935 | 0.101 | 0.543 | 0.897 |
|  | Epistatic effects | Uniform | 0.955 | 0.214 | 1.134 | 0.600 |
|  |  | Normal | 0.955 | 0.198 | 1.083 | 0.625 |
|  |  | Laplace | 0.947 | 0.192 | 1.064 | 0.633 |
| Weak epistasis<br>$w_j \sim \text{Unif}(0.5, 1.5)$,<br>$w_{jk} \sim \text{Normal}(1, 0.1^2)$ | Individual effects | Uniform | 0.920 | 0.126 | 0.589 | 0.819 |
|  |  | Normal | 0.915 | 0.119 | 0.564 | 0.838 |
|  |  | Laplace | 0.915 | 0.116 | 0.549 | 0.846 |
|  | Epistatic effects | Uniform | 0.952 | 0.195 | 1.140 | 0.237 |
|  |  | Normal | 0.957 | 0.161 | 1.081 | 0.260 |
|  |  | Laplace | 0.957 | 0.143 | 1.056 | 0.268 |

**Table 7:** Estimation performance with weak, moderate and strong epistatic interactions among 10 genetic features under different priors.

|  |  | Priors | Coverage | MAE | CI width | Correlation |
| --- | --- | --- | --- | --- | --- | --- |
| Strong epistasis<br>$w_j \sim \text{Unif}(0.5, 1.5)$<br>$w_{jk} \sim \text{Unif}(0.5, 1.5)$ | Individual effects | Uniform | 0.927 | 0.150 | 0.660 | 0.738 |
|  |  | Normal | 0.924 | 0.151 | 0.654 | 0.739 |
|  |  | Laplace | 0.917 | 0.154 | 0.641 | 0.738 |
|  | Epistatic effects | Uniform | 0.938 | 0.225 | 0.873 | 0.382 |
|  |  | Normal | 0.932 | 0.225 | 0.868 | 0.382 |
|  |  | Laplace | 0.922 | 0.226 | 0.857 | 0.383 |
| Moderate epistasis<br>$w_j \sim \text{Unif}(0.5, 1.5)$,<br>$w_{jk} \sim \text{Normal}(1, 0.3^2)$ | Individual effects | Uniform | 0.931 | 0.160 | 0.668 | 0.709 |
|  |  | Normal | 0.928 | 0.160 | 0.662 | 0.710 |
|  |  | Laplace | 0.918 | 0.162 | 0.649 | 0.711 |
|  | Epistatic effects | Uniform | 0.872 | 0.218 | 0.872 | 0.385 |
|  |  | Normal | 0.870 | 0.218 | 0.867 | 0.384 |
|  |  | Laplace | 0.865 | 0.217 | 0.856 | 0.387 |
| Weak epistasis<br>$w_j \sim \text{Unif}(0.5, 1.5)$,<br>$w_{jk} \sim \text{Normal}(1, 0.1^2)$ | Individual effects | Uniform | 0.910 | 0.142 | 0.637 | 0.809 |
|  |  | Normal | 0.909 | 0.143 | 0.630 | 0.810 |
|  |  | Laplace | 0.903 | 0.144 | 0.615 | 0.812 |
|  | Epistatic effects | Uniform | 0.947 | 0.112 | 0.835 | 0.214 |
|  |  | Normal | 0.947 | 0.109 | 0.829 | 0.217 |
|  |  | Laplace | 0.948 | 0.103 | 0.815 | 0.223 |

**Figure 4:**
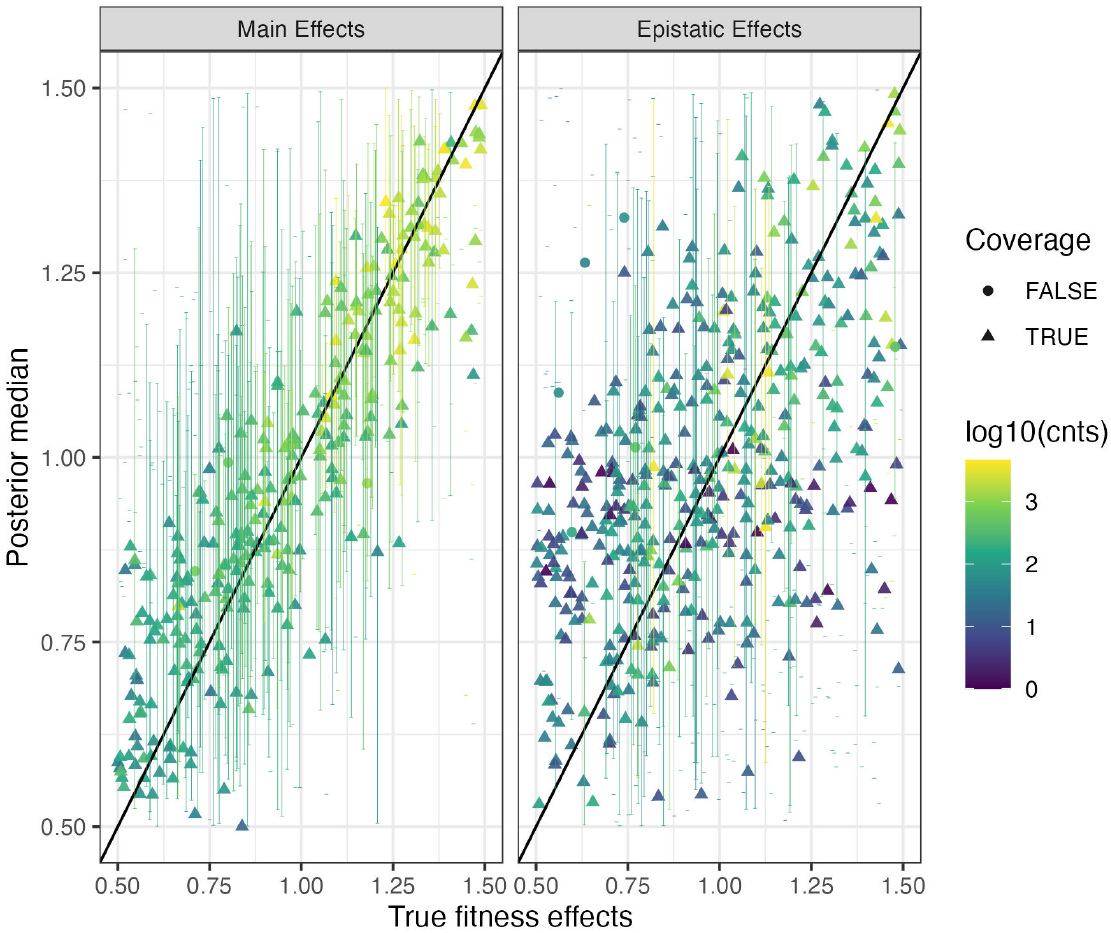
Estimation performance for the four-feature fitness model with strong epistasis using a Laplace prior, shown separately for main and epistatic effects. Posterior median estimates of mutational fitness effects are plotted against the true simulated fitness effects. Points are colored by the log-transformed number of occurrences of each mutation in the simulated alignment. Triangles indicate cases in which the true value falls within the 95% credible interval, whereas circles indicate cases in which it does not. Vertical error bars show 95% credible intervals, and the diagonal line denotes perfect agreement between estimated and true effects.

Finally, we evaluated the effect of omitting epistasis from the inference model when the true data-generating process included pairwise interactions. In the four-feature setting, the reduced (i.e misspecified) additive model performs poorly when epistatic effects are strong or moderate as coverage for individual effects decreases substantially and MAE increases relative to the full model (Table 8). Thus, ignoring even moderately strong epistasis can bias estimates of individual mutational effects. Under weak epistasis, however, the reduced model produces lower MAE and narrower intervals than the full model, although coverage also decreases. This result can be interpreted by considering the bias-variance trade-off in model complexity. If weak epistatic effects contribute noise rather than signal, including additional interaction terms increases model complexity without capturing meaningful effects, thereby degrading estimates of the individual additive effects.

**Table 8:** Estimation performance under a reduced model ignoring epistasis with weak, moderate and strong epistatic interactions of a 4-feature model.

|  |  | Priors | Coverage | MAE | CI width | Correlation |
| --- | --- | --- | --- | --- | --- | --- |
| Strong epistasis<br>$w_j \sim \text{Unif}(0.5, 1.5)$<br>$w_{jk} \sim \text{Unif}(0.5, 1.5)$ | Individual effects<br>(Full model) | Uniform | 0.938 | 0.108 | 0.556 | 0.885 |
|  |  | Normal | 0.933 | 0.107 | 0.537 | 0.890 |
|  |  | Laplace | 0.935 | 0.108 | 0.523 | 0.889 |
|  | Individual effects<br>(Reduced model) | Uniform | 0.720 | 0.127 | 0.377 | 0.848 |
|  |  | Normal | 0.720 | 0.127 | 0.373 | 0.847 |
|  |  | Laplace | 0.733 | 0.127 | 0.370 | 0.845 |
| Moderate epistasis<br>$w_j \sim \text{Unif}(0.5, 1.5)$,<br>$w_{jk} \sim \text{Normal}(1, 0.3^2)$ | Individual effects<br>(Full model) | Uniform | 0.943 | 0.105 | 0.575 | 0.889 |
|  |  | Normal | 0.940 | 0.101 | 0.556 | 0.896 |
|  |  | Laplace | 0.935 | 0.101 | 0.543 | 0.897 |
|  | Individual effects<br>(Reduced model) | Uniform | 0.695 | 0.130 | 0.370 | 0.821 |
|  |  | Normal | 0.705 | 0.129 | 0.366 | 0.822 |
|  |  | Laplace | 0.695 | 0.129 | 0.363 | 0.820 |
| Weak epistasis<br>$w_j \sim \text{Unif}(0.5, 1.5)$,<br>$w_{jk} \sim \text{Normal}(1, 0.1^2)$ | Individual effects<br>(Full model) | Uniform | 0.920 | 0.126 | 0.589 | 0.819 |
|  |  | Normal | 0.915 | 0.119 | 0.564 | 0.838 |
|  |  | Laplace | 0.915 | 0.116 | 0.549 | 0.846 |
|  | Individual effects<br>(Reduced model) | Uniform | 0.875 | 0.095 | 0.387 | 0.881 |
|  |  | Normal | 0.870 | 0.095 | 0.382 | 0.883 |
|  |  | Laplace | 0.863 | 0.096 | 0.378 | 0.885 |

### Application to QRDR mutations in *E. coli*

We applied our inference framework to investigate the fitness effects of quinolone resistance mutations in *E. coli* ST131 from a time-calibrated phylogeny with 1763 samples previously reconstructed from a core-genome single-nucleotide polymorphism (SNP) alignment (Figure 5A). We inferred fitness effects for common mutations in the quinolone resistance-determining region (QRDR) (Putten *et al*., 2019). These included resistance-associated mutations at Ser83 (*gyrA*-S83L) and Asp87 (*gyrA*-D87N) in *gyrA*; Ser80 (*parC*-S80I) and Glu84 (*parC*-E84V) in *parC*; and Ile529 (*parE*-I529L) in *parE* (Friedman *et al*., 2001; Onseedaeng and Ratthawongjirakul, 2016; Zakour *et al*., 2016; Johnson *et al*., 2015). Because *gyrA*-D87N and *parC*-S80I co-occurred in more than 95% of samples, we treated them as a single combined feature. Previous lab-based experiments demonstrated these QRDR mutations form an epistatic network of interacting sites with pairwise and higher-order interactions significantly impacting the fitness of genotypes with multiple QRDR mutations (Ben Zakour *et al*., 2016; Baker *et al*., 2013; Marcusson *et al*.,2009). We therefore inferred individual QRDR effects as well as first-order and second-order epistatic interactions under the fitness model in Eq. 4 using a Laplace shrinkage prior, which included a baseline fitness effect λ_0_ *∼* Unif(0.0, 2.0) and a branch-specific random term λ_*iϵ*_ *∼* log Normal 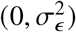 in the birth rate λ_*i*_ of branch *i* to account for fitness variation not captured by the QRDR features.

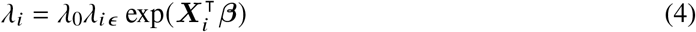

**Figure 5:**
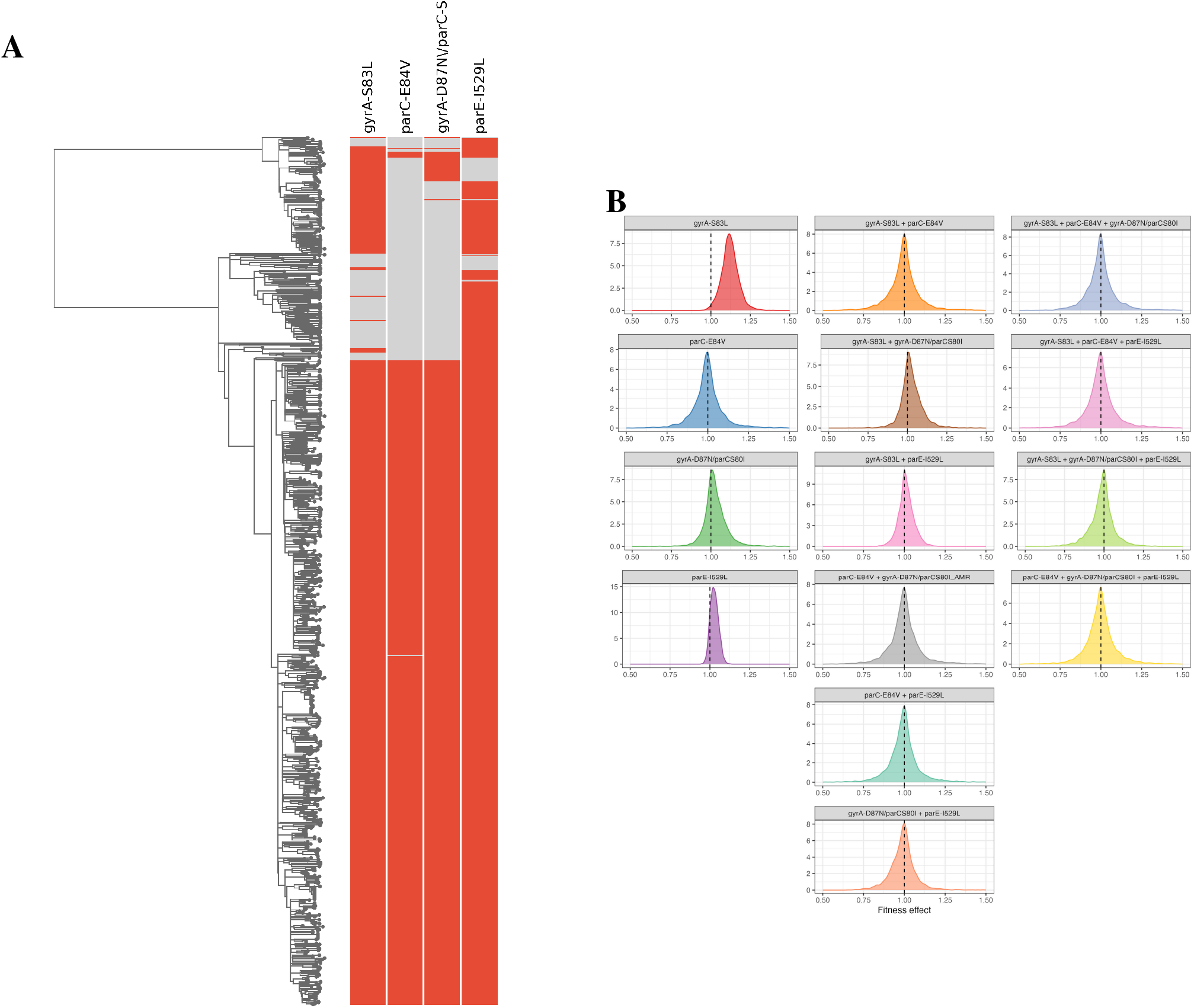
(A): Maximum-likelihood time-calibrated phylogeny of *E. coli* ST131. Feature presence in each sample is shown by colored bands: red indicates that the mutation is present, and grey indicates that the mutation is absent. (B) Posterior distributions of inferred fitness effects for QRDR mutations and epistatic interactions, ordered by the posterior median effect.

Most inferred individual fitness effects and epistatic interactions are centered near a neutral effect of one on the multiplicative fitness scale (Table 9; Figure 5B). Among the individual QRDR features, *gyrA*-S83L showed the greatest evidence of a positive fitness effect, with a posterior median of 1.118 and a 95% credible interval excluding 1.0, consistent with the previous findings of Kepler *et al*. (2024). The estimated effects of *parE*-I529L, *parC*-E84V, and the combined *gyrA*-D87N/*parC*-S80I feature are close to neutral (1.0), with credible intervals overlapping neutrality. First-order epistatic terms are also centered near 1.0, indicating limited evidence for strong pairwise epistasis among the QRDR mutations considered. Similarly, the second-order interaction terms have posterior medians near 1.0 and credible intervals spanning neutrality, providing little evidence for higher-order epistasis. Overall, these results indicate that the primary QRDR-associated fitness signal in this ST131 dataset is the positive effect of *gyrA*-S83L, with little evidence for first- and second-order epistasis among QRDR mutations.

**Table 9:** Estimated fitness effects and credible intervals for individual effects, first-order epistasic, and second-order epistasic interactions in *E. coli* ST131.

|  | Feature | Number of occurrences | est_med | CI_L | CI_U |
| --- | --- | --- | --- | --- | --- |
| Individual effects | gyrA-S83L | 1547 | 1.118 | 1.013 | 1.228 |
|  | parC-E84V | 1323 | 0.995 | 0.813 | 1.188 |
|  | gyrA-D87N/parC-S80I | 1372 | 1.019 | 0.906 | 1.186 |
|  | parE-I529L | 1684 | 1.026 | 0.982 | 1.082 |
| Pairwise epistasis | gyrA-S83L + parC-E84V | 1323 | 0.995 | 0.816 | 1.182 |
|  | gyrA-S83L + gyrA-D87N/parC-S80I | 1372 | 1.018 | 0.888 | 1.178 |
|  | gyrA-S83L + parE-I529L | 1494 | 1.009 | 0.919 | 1.117 |
|  | parC-E84V + gyrA-D87N/parC-S80I | 1323 | 0.998 | 0.830 | 1.240 |
|  | parC-E84V + parE-I529L | 1323 | 0.996 | 0.821 | 1.193 |
|  | gyrA-D87N/parC-S80I + parE-I529L | 1324 | 0.993 | 0.821 | 1.150 |
| Three-way epistasis | gyrA-S83L + parC-E84V + gyrA-D87N/parC-S80I | 1323 | 0.995 | 0.817 | 1.180 |
|  | gyrA-S83L + parC-E84V + parE-I529L | 1323 | 0.996 | 0.814 | 1.202 |
|  | gyrA-S83L + gyrA-D87N/parC-S80I + parE-I529L | 1324 | 0.991 | 0.829 | 1.166 |
|  | parC-E84V + gyrA-D87N/parC-S80I + parE-I529L | 1323 | 0.998 | 0.829 | 1.184 |

## Discussion

Learning how genotype maps to fitness is essential for understanding what genomic features (mutations, genes, mobile genetic elements) shape the fitness and evolutionary potential of pathogens. While lab-based experimental studies can accurately determine the fitness effects of individual variants, it is generally unclear how fitness measured in the lab translates to population-level fitness and thus epidemic potential in real-world settings. In addition, epistasis can restrict accessible evolutionary paths and make adaptation depend on genetic background (Weinreich *et al*., 2006; Martin *et al*., 2007; Flynn *et al*., 2013). Here, we developed a gradient-based Bayesian MFBD framework for estimating additive and epistatic genotype-to-fitness maps from pathogen phylogenies. This framework builds on phylodynamic birth-death models that connect branching patterns in a phylogeny to lineage-specific transmission or growth rates (Volz *et al*., 2013; Barido-Sottani *et al*., 2020; Stadler and Bonhoeffer, 2013; Rasmussen and Stadler, 2019). Compared with maximum-likelihood MFBD approaches for multi-feature fitness inference (Kepler *et al*., 2021, 2024), the Bayesian formulation provides full posterior uncertainty for mutational and epistatic fitness effects and allows shrinkage priors to regularize weakly supported terms.

A gradient-based Bayesian implementation like HMC is especially useful for this class of genotype-to-fitness models because the posterior can be high-dimensional, correlated, and weakly identified for rare features or interaction terms (Betancourt, 2018). Previous phylodynamic work has used HMC to improve Bayesian inference for coalescent-based population dynamics and episodic birth-death-sampling models, showing that gradient-based samplers can substantially improve efficiency over conventional MCMC approaches (Lan *et al*., 2015; Shao *et al*., 2024). In addition, the phylodynamic likelihood (Eq. 1) assumes that birth, death, and sampling events are all conditionally independent between lineages given the model parameters. We exploit this model structure to vectorize phylogenetic trees into arrays, allowing the likelihood and its gradients to be evaluated in parallel. As a result, the framework improves computational efficiency, while preserving posterior uncertainty in poorly supported regions of the genotype-to-fitness map.

The simulation study shows that additive fitness effects can be estimated accurately with fairly low levels of uncertainty when the phylogeny contains sufficient information. In both the four-feature and ten-feature additive settings, Bayesian estimates are comparable to maximum-likelihood estimates, and empirical accuracy improves as tree size increases (Tables 1, 3; Figures 2, 3). This pattern is consistent with the basic premise of fitness-dependent phylodynamics, where repeated occurrences of a feature, together with the branching history of lineages that carry it, provide information about its population-level fitness effect (Rasmussen and Stadler, 2019; Kepler *et al*., 2021). One exception is that the fitness effects of strongly deleterious mutations tend to be biased upward, making them appear less harmful than their true effects (Figure 2A,Figure 3A,Figure 4). This bias is expected because deleterious mutations are more likely to occur on lineages that die out quickly, leaving few descendants and few opportunities for sampling. The deleterious mutations that remain in the reconstructed tree are therefore a nonrandom subset, often carried by lineages that persist because of chance, favorable genetic backgrounds, or favorable epidemiological conditions. Finally, the higher MAE in the ten-feature setting indicates that phylogenetic information becomes diluted as the number of estimated effects increases relative to the number of informative lineages. Thus, our method’s performance is driven less by the total number of features than by how often each feature appears independently across the phylogeny.

The results also clarify when imposing regularization through shrinkage priors is useful. In correctly specified additive models, shrinkage priors do not improve accuracy over the weakly informative uniform priors, suggesting limited benefit when few parameters are estimated from informative data. By contrast, shrinkage improves estimates for epistatic terms with small true fitness effects, especially in the four-feature simulations where Laplace and normal priors reduced MAE and interval width (Table 6), reflecting a bias-variance trade-off. Shrinkage can stabilize weak interaction estimates, but it cannot compensate for absent genotype combinations. The reduced-model simulations show the complementary risk, since omitting moderate or strong epistasis could bias individual effects, whereas fitting weak unsupported interactions increases uncertainty and reduces coverage (Table 8). Model choice should therefore depend on both biological expectations about epistasis and the co-occurrence information available in the phylogeny. In summary, larger phylogenies improve the accuracy of fitness-effect estimation under the MFBD model. For additive landscapes without epistasis, shrinkage priors does not materially improve accuracy relative to a flat prior. When pairwise epistasis is included, interaction effects are less precisely estimated than individual mutational effects, but increasing the number of interaction terms does not significantly reduce overall accuracy.

Applying our framework to the time-calibrated phylogeny of *E. coli* ST131 identifies *gyrA*-S83L as the QRDR mutation with the strongest evidence for a positive population-level fitness effect. This finding is consistent with Kepler *et al*. (2024), who also identified *gyrA*-S83L as a major determinant of ST131 fitness. Laboratory studies suggest several mechanisms by which the *gyrA*-S83L mutation could be beneficial, including altered DNA supercoiling and stress-response gene expression (Marcusson *et al*., 2009; Baker *et al*., 2013; Webber *et al*., 2013). Other work suggests that quinolone-resistance backgrounds may also affect the carriage or stability of IncF plasmids, which are common in ST131 and often carry antimicrobial resistance (Toukdarian, 2004; Villa *et al*., 2010; Pitout and Finn, 2020). Our analysis extends previous work by considering a broader set of QRDR features and by explicitly estimating possible epistasic interactions among them. The estimated fitness effects of two-way and three-way interaction terms between individual mutations are centered near neutrality with credible intervals spanning one. This suggests that idiosyncratic epistasis does not strongly modify the large beneficial effect of *gyrA*-S83L across different QRDR genotypes. Previous experimental work has shown epistatic interactions between certain beneficial mutations (He *et al*., 2010; Khan *et al*., 2011) and between antimicrobial-resistance mutations (Remold and Lenski, 2004; Trindade *et al*., 2009) in *E. coli*. Further, evidence has shown that epistatic interactions exist among QRDR mutations in both *Salmonella* and in *E. coli* (Marcusson *et al*., 2009; Baker *et al*., 2013). However, our phylodynamic inferences provide limited evidence that QRDR interactions explain additional variation in fitness. Several factors may explain this discrepancy. First, many QRDR mutations co-occur on similar genetic backgrounds, reducing our power to separate the effects of different interactions from main effects. Second, MFBD models infer lineage fitness from sampled pathogen phylogenies, which reflect transmission, sampling, and lineage expansion in the host population, whereas laboratory studies usually measure a single component of fitness such as growth or resistance under controlled experimental conditions (Hinz *et al*., 2024). It could therefore be the case that epistasis strongly impacts individual components of fitness, but that these epistatic effects are weaker for the higher-level and more integrative measures of fitness obtained from phylogenies. Finally, some apparent epistasis may arise because mutations act additively on an intermediate phenotype, such as protein stability, enzyme activity, drug resistance, or replication rate, while fitness is a nonlinear function of that phenotype (Starr and Thornton, 2016; Otwinowski *et al*., 2018; Diaz-Colunga *et al*., 2023).

Several limitations define the current scope of inference and point to useful extensions. The simulations assumed a fixed phylogeny, reconstructed lineage genotypes, and known fixed parameters, whereas empirical applications should account for uncertainty in tree topology, node times, ancestral states, sampling processes and background epidemiological heterogeneity (Drummond and Rambaut, 2007; Hanson-Smith *et al*., 2010). In the ST131 analysis, the branch-specific random effect absorbed fitness variation not captured by QRDR mutations, but it did not identify the genomic or ecological sources of that residual variation, which may include other resistance genes, plasmids, virulence factors, host populations, geography, and treatment environments (Zakour *et al*., 2016; Kepler *et al*., 2024; Fuzi and Sokurenko, 2023). Future work should therefore propagate phylogenetic and ancestral-state uncertainty, incorporate richer genomic and epidemiological covariates, and use model-selection or hierarchical shrinkage strategies to balance epistatic complexity against available co-occurrence information (Park and Casella, 2008; Carvalho *et al*., 2010). More broadly, phylodynamic genotype-to-fitness inference complements experimental fitness-landscape approaches. Controlled lab-based experiments can measure fitness effects precisely but may depend on assay conditions and scale poorly to large combinatorial spaces, while phylodynamic models estimate population-level fitness effects from sampled pathogen phylogenies and make explicit when the data do, or do not, support a genetic association with lineage fitness (Fowler and Fields, 2014; Blanquart and Bataillon, 2016; Zhou *et al*., 2022; Weinreich *et al*., 2006; Flynn *et al*., 2013).

## Conclusion

In conclusion, we developed a gradient-based Bayesian MFBD framework for estimating additive and epistatic genotype-to-fitness maps from pathogen phylogenies. Across simulations, the framework recovers additive fitness effects with proper quantification of uncertainty and shows that inference improves most consistently when phylogenies contain enough independent information for each feature. Epistatic effects are harder to estimate because they depend on mutation co-occurrence, but shrinkage priors help regularize weakly supported interaction terms. In the *E. coli* ST131 application, *gyrA*-S83L shows the strongest evidence for a positive population-level fitness effect, whereas QRDR interaction terms are centered near neutrality. These results show that Bayesian phylodynamic inference can identify well-supported genetic contributors to pathogen fitness in real-world environments while explicitly quantifying uncertainty

## Acknowledgements

D.A.R. and S.C. were partially supported by the U.S. National Science Foundation award 2142931.

## Data availability statement

All project code, documentations and data are available at https://github.com/ShiCenNCSU/BayesMFBD.

